# *ComBat-Seq*: batch effect adjustment for RNA-Seq count data

**DOI:** 10.1101/2020.01.13.904730

**Authors:** Yuqing Zhang, Giovanni Parmigiani, W. Evan Johnson

## Abstract

The benefit of integrating batches of genomic data to increase statistical power in differential expression is often hindered by batch effects, or unwanted variation in data caused by differences in technical factors across batches. It is therefore critical to effectively address batch effects in genomic data. Many existing methods for batch effect adjustment assume continuous, bell-shaped Gaussian distributions for data. However in RNA-Seq studies where data are skewed, over-dispersed counts, this assumption is not appropriate and may lead to erroneous results. Negative binomial regression models have been used to better capture the properties of counts. We developed a batch correction method, ComBat-Seq, using negative binomial regression. ComBat-Seq retains the integer nature of count data in RNA-Seq studies, making the batch adjusted data compatible with common differential expression software packages that require integer counts. We show in realistic simulations that the ComBat-Seq adjusted data result in better statistical power and control of false positives in differential expression, compared to data adjusted by the other available methods. We further demonstrated in a real data example where ComBat-Seq successfully removes batch effects and recovers the biological signal in the data.

## Introduction

Genomic data are often produced in batches due to logistical or practical restrictions, but technical variation and differences across batches, often called batch effects, can cause significant heterogeneity across batches of data (Leek *et al.*, 2010). Batch effects often result in discrepancies in the statistical distributions across data from different technical processing batches, and can have unfavorable impact on downstream biological analysis. The presence of batch effects often reduces the benefits of integrating batches of data to increase the inferential power to discover relevant biology from the combined data.

Batch effects often cannot be fully addressed by normalization methods and procedures. The differences in the overall expression distribution of each sample across batch may be corrected by normalization methods, such as transforming the raw counts to (logarithms of) CPM, TPM, or RPKM/FPKM, the trimmed mean of M values (TMM, Robinson and Oshlack, 2010), or relative log expression (RLE, Risso *et al.*, 2014a). However, batch effects in composition, i.e. the level of expression of genes scaled by the total expression (coverage) in each sample, cannot be fully corrected with normalization. Leek *et al.* (2010) provided an example of composition batch effects in microarray data, showing that while the overall distribution of samples may be normalized to the same level across batches, individual genes may still be affected by batch-level bias.

Many methods have been proposed to address batch effects in RNA-Seq studies. For example, ComBat (Johnson *et al.*, 2007) remains one of the most popular batch effect adjustment methods when the effects come from known sources. For heterogeneity from unknown sources, SVASeq (Leek, 2014) and RUVSeq (Risso *et al.*, 2014b) are commonly used. Methods designed for specific downstream tasks have also been proposed, including our own work using reference batches for biomarker development and training (Zhang *et al.*, 2018). For differential expression, many common methods or procedures (e.g. edgeR (Robinson *et al.*, 2010) and DESeq2 (Love *et al.*, 2014)) suggest to include batch variables as covariates in the linear models behind these methods to account for the impact of batch.

Despite the established progress, there are still gaps in batch adjustment methodology for RNA-Seq data which need to be bridged. Often times, batch effect adjustment methods either do not directly provide adjusted data with batch effects removed, or do not preserve the integer nature of counts in the adjusted data, despite the requirement of software such as edgeR and DESeq2 that specifically require untransformed count matrices as inputs. This results in an inconsistency in the analysis pipeline of RNA-Seq studies, as batch corrected data cannot be used as inputs for these differential expression software. For this practical issue, it is favorable to develop a method which generates adjusted data and is able to preserve the count nature of data.

More importantly, many popular adjustment methods, including ComBat, assume Gaussian distributions for the underlying distribution of the data, which is not an appropriate distributional assumption for counts. These methods typically estimate parameters representing differences in the statistical moments across batches (usually the mean and the variance). Then they adjust all batches to the same overall level in these moments. Such adjustment does not preserve integers, and may results in negative values in adjusted count matrix, which is difficult to interpret biologically. In addition, it has been well-established that there exists a mean-variance dependence in RNA-Seq count data (Law *et al.*, 2014). Distributions of counts are skewed and over-dispersed, i.e. the variance is often larger than the mean of gene expression and genes with smaller counts tend to have larger variances. These properties cannot be reflected with Gaussian distribution, which assumes independent mean and variance parameters. Negative binomial regression models have been widely used to model count data in RNA-Seq studies. The Negative binomial distribution has the potential to describe the skewness and mean-variance relationship observed in count matrices. We propose to extend the original ComBat framework to RNA-Seq studies using negative binomial regression.

Finally, existing methods may not be flexible enough to address all types of batch effects. In particular, including batch variables in software for differential expression may be sufficient to account for batch effects in the mean expression. However, since both software assume a single dispersion parameter for all samples, variance batch effects is restricted, and completely determined by mean batch effects, due to the properties of negative binomial modeling. Such assumption is strong and may not always hold for real data. Therefore, we propose a more flexible approach to address batch effects in the variance.

In this paper, we present a batch effect adjustment method, ComBat-Seq, that extends the original ComBat adjustment framework to address the challenges in batch correction in RNA-Seq count data. It generates adjusted data in the form of counts, thus preserving the integer nature of data. We demonstrate that ComBat-Seq adjustment has potential benefits in differential expression compared to the other adjustment methods, especially when there is a large variance batch effect in the data.

## Materials and methods

We propose a negative binomial regression model to estimate batch effects based on the count matrix in RNA-Seq studies. With the estimated batch effect parameters, we calculate “batch-free” distributions, i.e. the expected distributions if there were no batch effects in the data. We then adjust the data by mapping the quantiles of the empirical distributions of data to the batch-free distributions. We further describe the methods in the sections below.

### ComBat-Seq model

We define a regression model for each gene. Let the expression count value for gene *g* of sample *j* from batch *i* be denoted by *y*_*gij*_. We assume that *y*_*gij*_ follows a negative binomial distribution *NB*(*µ*_*gij*_, *ϕ*_*gi*_), where *µ*_*gij*_ and *ϕ*_*gi*_ are the mean and the dispersion parameters. We propose the gene-wise model:

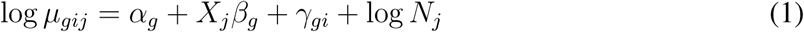

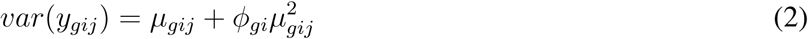

where *α*_*g*_ denotes the logarithm of expected counts for “negative” samples. *X*_*j*_*β*_*g*_ reflects changes to the log of expected counts due to biological conditions, which is preserved in the data after adjustment. In this term, *X*_*j*_ may be an indicator of the biological condition for sample *j*, or a continuous value for a clinical covariate. *β*_*g*_ denotes the corresponding regression coefficient. *N*_*j*_ represents the library size, i.e. total counts across all genes in sample *j*. The mean and dispersion batch effect parameters are denoted by *γ*_*gi*_ and *ϕ*_*gi*_, respectively, modeling the effect of batch *i* on gene *g*. We estimate the model parameters, especially batch effect parameters *γ*_*gi*_ and *ϕ*_*gi*_, following the established methods in edgeR (Robinson *et al.*, 2010; McCarthy *et al.*, 2012; Chen *et al.*, 2014). Specifically, the mean batch effect parameters *γ*_*gi*_s are estimated with Fisher scoring iteration, implemented in an optimized way to reduce the computational time. The dispersion parameters *ϕ*_*gi*_ are estimated gene-wise by maximizing the Cox–Reid adjusted profile likelihood (APL), as defined in Chen *et al.* (2014), and results in non-negative dispersion estimates. Note that the estimates for mean of expression are not required to be non-negative, since they are on the log scale. We estimated the gene-wise dispersion within each batch in ComBat-Seq.

### ComBat-Seq adjustment

After the modeling, we obtain estimated batch effect parameters 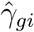 and 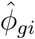, as well as the fitted expectation of count 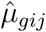. We then calculate parameters for “batch-free” distributions as follows: we assume that the adjusted data 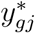 follows a “batch-free” negative binomial distribution 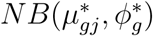, where parameters are calculated as

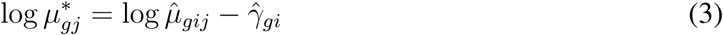

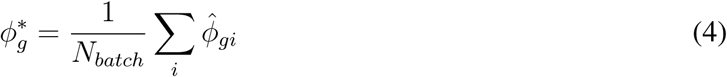

Then, the adjusted data 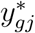 is calculated by finding the closest quantile on the batch-free distribution to the quantile of the original data *y*_*gij*_ on the empirical distribution, estimated as 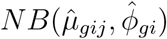. Specifically, we find the adjusted value 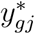 such that 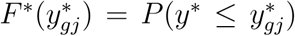 is closest in absolute value to *F* (*y*_*gij*_) = *P* (*y ≤ y*_*gij*_). Zero counts are mapped to zeros. We perform this mapping for every value in the count matrix, which completes the adjustment. Figure 1 summarizes the whole workflow of ComBat-Seq.

**Figure 1:**
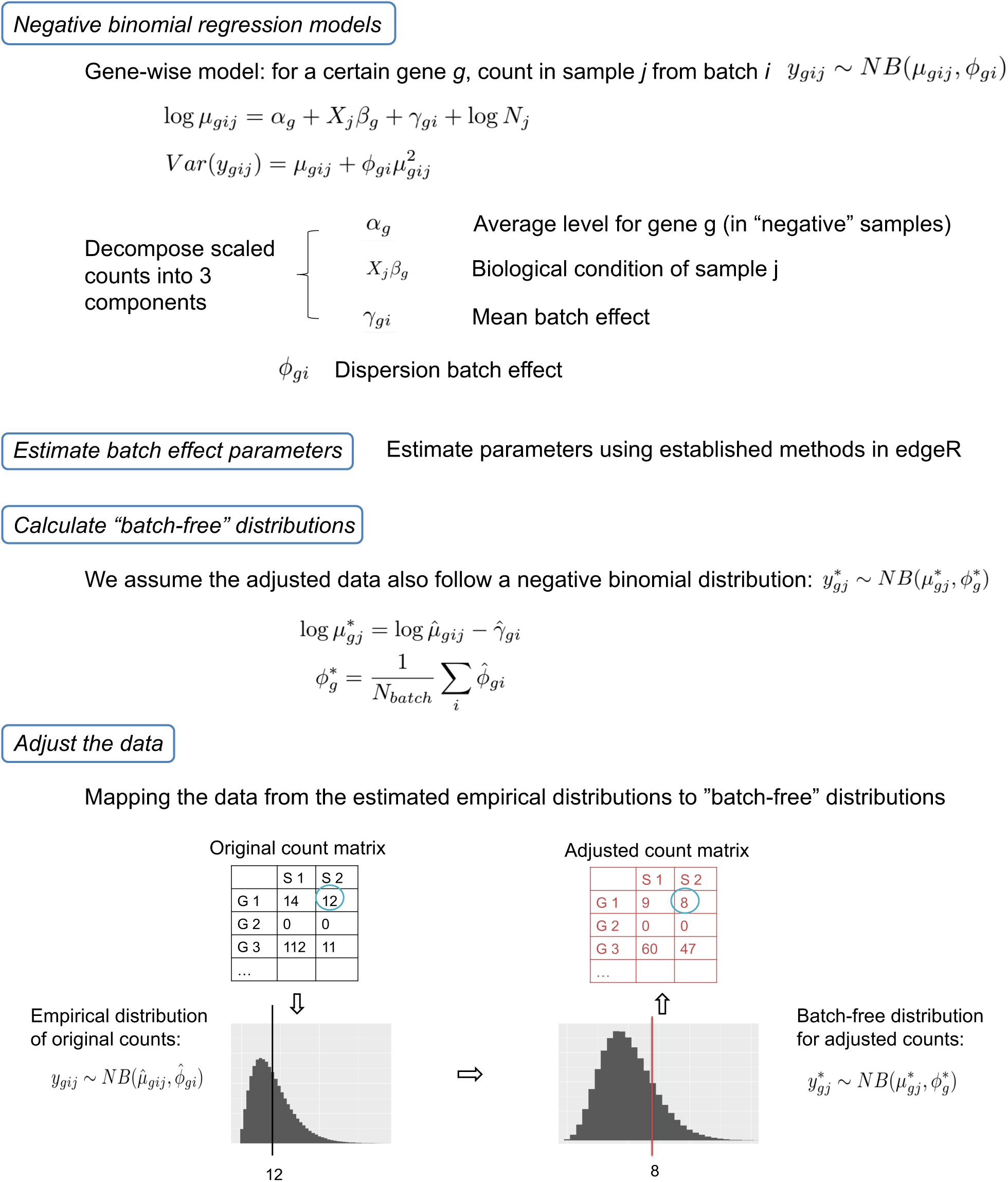
A diagram for the ComBat-Seq modeling and adjustment workflow.

### Simulations

We evaluated the performance of ComBat-Seq with simulation experiments consisting of three steps: 1) we simulated RNA-Seq studies with biological conditions and batch effects, 2) adjusted the batch differences with ComBat-Seq as well as other available methods, and 3) evaluated the performance of batch effect adjustment by the impact on differential expression using the adjusted data.

We used the *polyester* R package (Frazee *et al.*, 2015) to simulate realistic RNA-Seq studies, which are in the form of gene-by-sample count matrices. We designed two biological conditions and two batches of samples. The *polyester* package human genome reference example provides information for 918 genes which we divided them into two groups: group 1 has higher expression in batch 2 and lower in batch 1, while group 2 has the reversed pattern, higher in batch 1 and lower in batch 2. This forms a batch effect in the “composition” of expression as described in the Introduction section, which cannot be fully addressed by normalization. We assume a biological variable with two levels, “negative (0)” and “positive (1)”, which can represent “control” and “tumor” samples in a real dataset, for example. We simulated both up-regulated and down-regulated true differentially expressed genes in both gene groups with increased expression in the positive (up-regulated) or the negative (down-regulated) biological condition. The remaining genes are only affected by batch, not by the condition. Differences in the average count of genes are simulated by specifying fold changes across biological and batch sample groups using *polyester*. We also made the dispersion of batch 2 a number of times larger than that of batch 1, allowing for different true dispersion parameters across batches. Figure 2 shows the design we used for simulated data.

**Figure 2:**
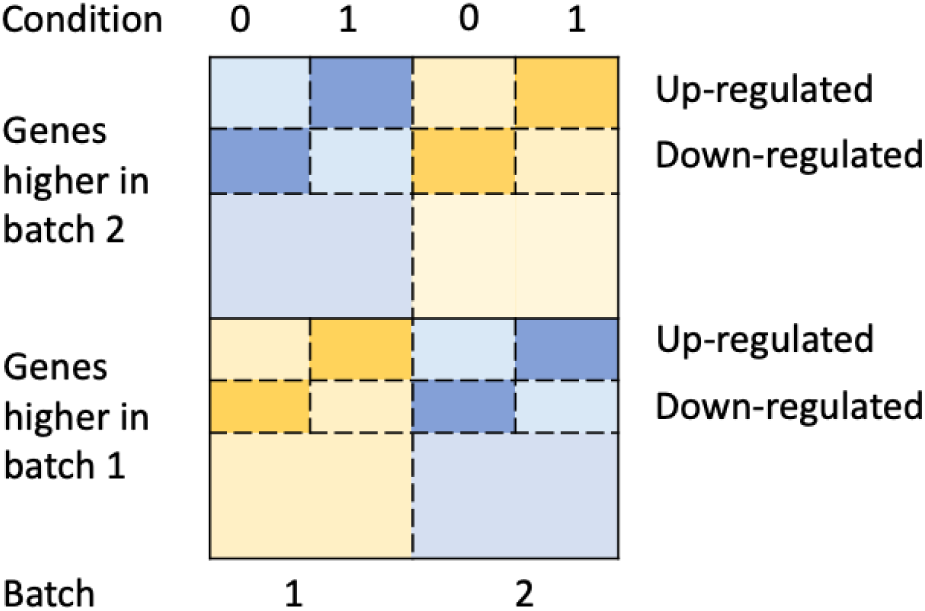
Study design of the simulation experiments. The data are in the form of gene by sample matrices. We simulated 2 biological conditions: negative (0) and positive (1), as well as 2 batches. We also simulated 2 groups of genes. Group 1 has higher expression in batch 2 and lower in batch 1, while group 2 has the reversed pattern, higher in batch 1 and lower in batch 2. The figure shows high expression in yellow, and low expression in blue. We also simulated differentially expressed genes in both groups. A deeper color represents increased expression due to biological condition.

We repeated the simulation while varying the level of mean and dispersion batch differences. Specifically, we changed the parameters for simulation such that mean of batch 2 is 1.5, 2, or 3 times that of batch 1. The dispersion of batch 2 was set to be 2, 3 or 4 fold of that of batch 1. Experiments with no mean batch effect or no dispersion differences were also included. Results were averaged over 100 repeated simulations under each parameter setting.

Our selected parameters in simulations are consistent with the degree of batch effect in real data, as summarized in Table 1. Our observed condition signal (fold change in expression) from real studies range from 1.65 to 3.98, and we specified a biological signal of 2 fold in simulations. With regard to batch differences in the moments, we observed mean batch effect to be in the range of 1.62 and 1.88 fold, and variance difference to be in 1.26 to 7.09 fold. Our selected parameters in the simulations align with the realistic range, suggesting that the results are likely representative for the expected effect on real data batch adjustment.

**Table 1:** Levels of biological and batch effects in real datasets. For the condition effect, we used edgeR to perform differential expression within one of the batches in the studies, and identified the top-50 up-regulated, and top-50 down-regulated genes, ranked by FDR corrected P values. We took the median of fold changes across conditions among the top-50 up-regulated genes, and the median of those among the down-regulated genes. We reported the maximum of two medians in the table. For mean batch effects, we calculated the gene-wise average expression within each batch, and each biological condition. We took the median of mean expression across genes, then compared the medians across batch, and report the maximum fold change. For dispersion differences, we report the maximum fold change in the median gene-wise dispersion across batch. Gene-wise dispersion are estimated with edgeR.

| Dataset | Condition | Batch Mean | Batch Dispersion | References |
| --- | --- | --- | --- | --- |
| GFRN | 1.65 | 1.88 | 2.78 | Rahman *et al.* (2017) |
| CHD8 | 2.92 | 1.68 | 7.09 | Sugathan *et al.* (2014) |
| TB | 3.98 | 1.62 | 1.26 | Zak *et al.* (2016); Suliman *et al.* (2018) |

The batch effects in mean and dispersion (variance) were adjusted with ComBat-Seq, the “one-step” approach, i.e. to include the batch variable in differential expression linear models, as well as with SVA-Seq and RUV-Seq. We also included another commonly used method in practice, which is to transform the count matrix to logCPM, then use the batch correction methods designed for Gaussian distributed data, such as the original ComBat method. Aside from batch adjusted data, we included two additional experiments for comparison: differential expression performed on 1) data without simulated batch effects, and 2) data with simulated batch differences, but no adjustment. We compared both the statistical power (true positive rate, TPR) and control of type-I errors (false positive rate, FPR) in detection using data without batch effects, data with batch effects before and after different adjustments.

### Real data application

We applied the proposed ComBat-Seq approach on an RNA-Seq data from a perturbation experiment using primary breast tissue attempting to profile the activity levels of growth factor receptor network (GFRN) pathways in relation to breast cancer progression (Rahman *et al.* (2017); McQuerry *et al.* (2019)). We took a subset of experiments, which consists of 3 batches. In each batch, the expression of a specific GFRN oncogene was induced by transfection to activate the down-stream pathway signals (different oncogene/pathway in each batch). Controls were transfected with a vector that expresses a green fluorescent protein (GFP), and GFP controls were present in all batches. More specifically, batch 1 contains 5 replicates of cells overexpressing HER2, and 12 replicates for GFP controls (GEO accession GSE83083); batch 2 contains 6 replicates of each for EGFR and its corresponding controls (GEO accession GSE59765); batch 3 consists of 9 replicates of each for wild type KRAS and GFP controls (GEO accession GSE83083).

Note that this is a challenging study design for batch effect adjustment: the control samples are balanced across batches, while each of the 3 kinds of treated cells, with different levels of biological signals, is completely nested within a single batch. A favorable adjustment would pool control samples from the three batches, while keeping all treated cells separated from the controls, and from each other.

We combined the three batches and performed batch correction. Among the batch correction methods considered, only RUV-Seq, the original ComBat used on logged and normalized data, and ComBat-Seq output adjusted data. We apply these methods to address the batch effects in the pathway signature dataset. We compared ComBat-Seq with the other methods, both qualitatively through principal component analysis (PCA), and quantitatively with explained variations by condition and batch.

## Results

We developed and implemented ComBat-Seq as described above, and applied the method to the simulated and real data examples. In the sections below, we justify the necessity of using negative binomial distribution instead of Gaussian distribution for count data. We then summarize our observations in simulations and a real data application example, showing the potential benefits of ComBat-Seq adjustment.

### Using appropriate model assumptions for count data

An example demonstrating the weakness of Gaussian-based models is given in Figure 3. In this example, we simulated a count matrix using *polyester* (Frazee *et al.*, 2015) with balanced case-control design and 2 batches. Figure 3 shows the counts in a gene. The control samples in both batches have low expression, while there is a case sample in batch 2 with a relatively large count over 30. When estimating the differences in mean across batch, due to the sample with the large count in the second batch, the mean of the second batch is estimated to be larger than that of batch If we apply Gaussian-based batch adjustment which brings the mean to the same level, control samples in the second batch will be adjusted to negative values, while counts in the first batch will be increased. This results in a significant artificial difference between control samples from the two batch after correction (*P* = 0.0033). These observations demonstrate the potential issue of applying batch correction method using Gaussian distribution on count data. A more appropriate model for integer counts would avoid such limitations.

**Figure 3:**
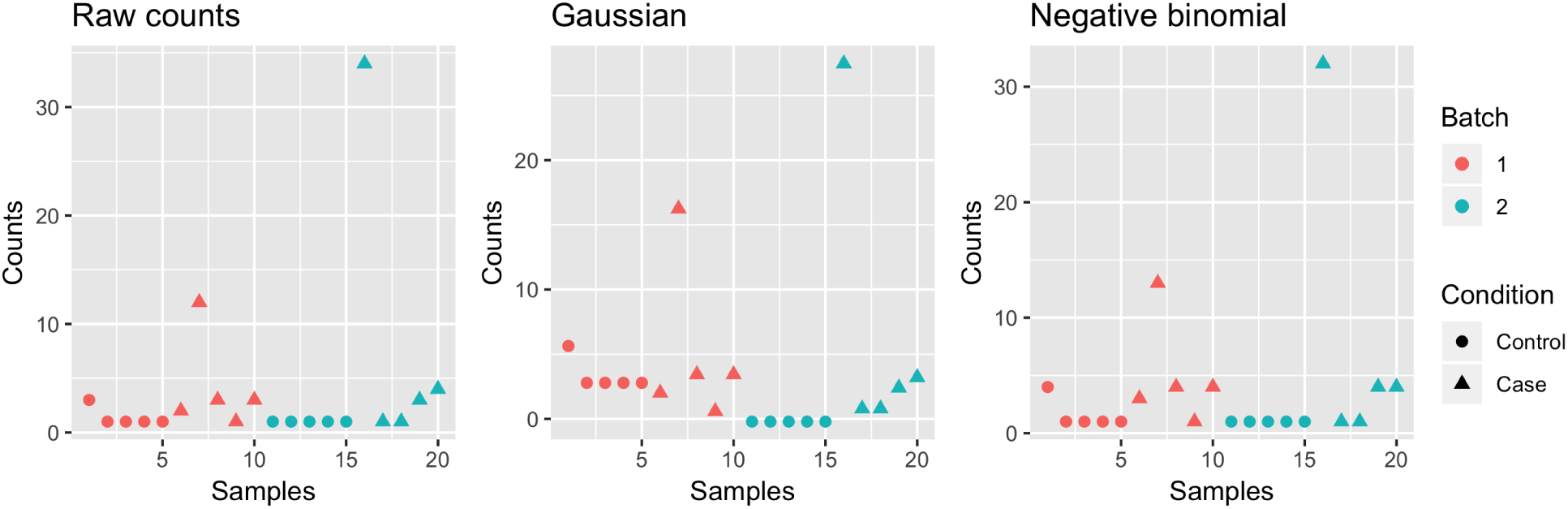
Problematic results caused by applying Gaussian-based batch adjustment approach on count data. We simulated count matrix with a balanced case-control design and 2 batches. The figure shows counts in a gene which is lowly expressed in almost all control samples. However, one case sample in the second batch contains a high count. Adjustment based on Gaussian distribution brings the mean of two batches to the same level, causing artificially induced differences across control samples from the two batches. When applying ComBat-Seq which is based on negative binomial distribution, the adjusted data no longer contain the negative values or the erroneous significant difference between control samples from the two batches.

We then applied the ComBat-Seq method, which assumes negative binomial distributions for the underlying data. As shown in Figure 3, the adjusted data do not contain negative values or the false significant result between the control samples of the two batches. This suggest that the negative binomial assumption indeed addresses the limitations mentioned above.

### Simulations

We evaluated ComBat-Seq and compared its performance with other available approaches in our simulation studies as described in the Method section. Results comparing all batch adjustment methods under different settings of degree of batch effects are summarized in Figure 4.

**Figure 4:**
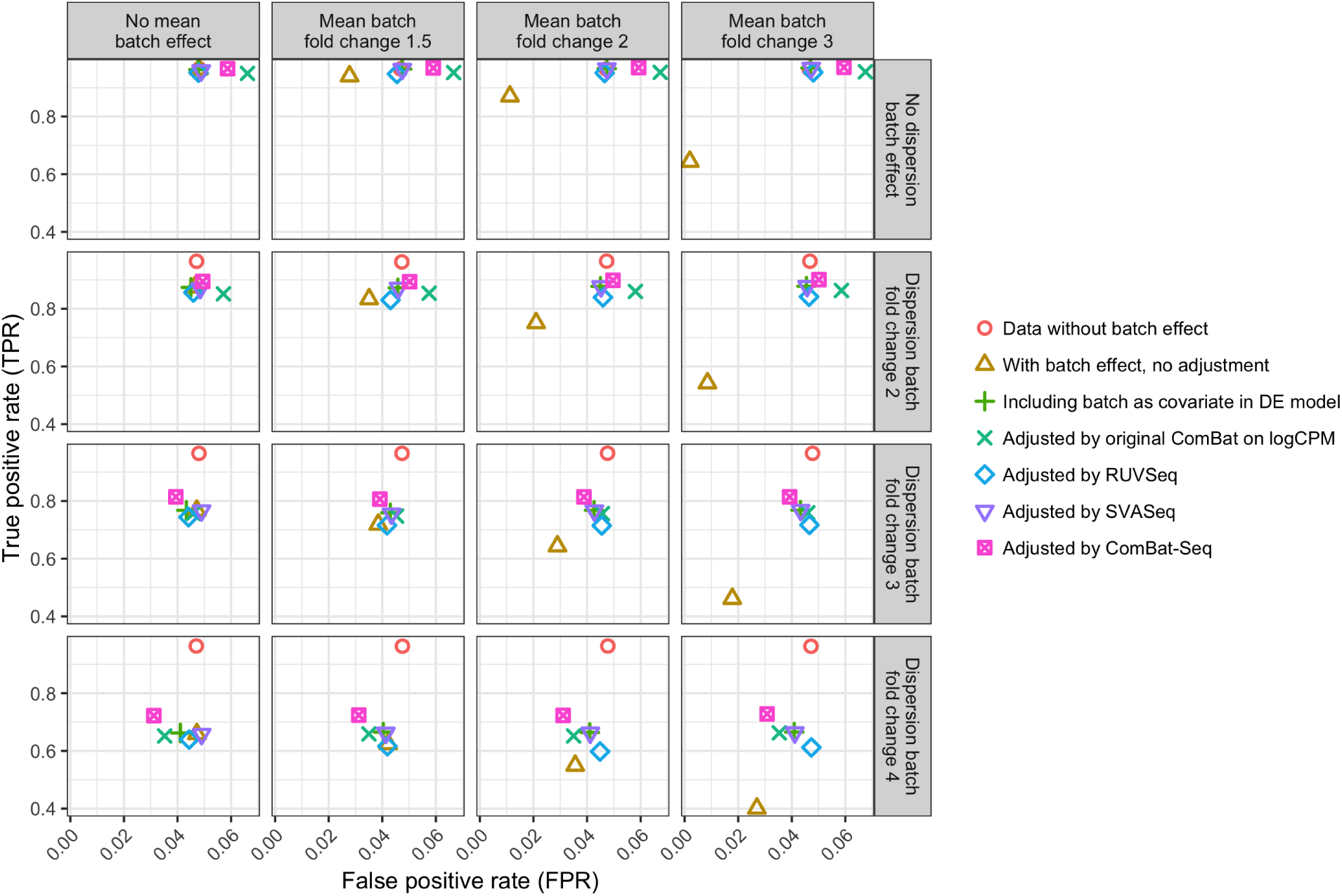
Simulation results under increasing level of differences across batch in the mean and variance of expression. Batch effects in the mean or the variance will cause a loss of power for differential expression detection. While all methods are able to increase the power for analysis, ComBat-Seq generally achieves the best power. Also, when there is a sufficient level of dispersion differences across batch, ComBat-Seq is able to better control false positives than the other methods.

Having batch effects in the data causes decrease in both true and false positive rates, compared to data without batch effects (FPR: 0.048, TPR: 0.96). For example, having only batch effects in the mean of at 1.5 fold but no dispersion differences reduces FPR to 0.028, and TPR to 0.94. Having only a 2 fold dispersion batch effect results in a 0.046 FPR and a 0.88 TPR. A larger difference in the mean or dispersion leads to a larger decrease in the power of detection in the unadjusted data. For instance, data with only mean batch effect but at a 3-fold difference have a 0.64 TPR, and data with no mean batch effect but a 4-fold dispersion effect have a 0.66 TPR (Figure 4). All batch correction methods are able to improve power for detection after adjusting the data with batch effects with increasing benefit as the degree of batch differences becomes more severe. ComBat-Seq achieves the highest true positive rate in general compared to the other batch correction methods. In a realistic range of a 1.5-fold mean batch effect, and a 2-fold dispersion batch effect, we observed a 0.89 TPR from ComBat-Seq, which is higher than the other methods (including batch as a covariate: 0.87, original ComBat on logCPM: 0.85, RUV-Seq: 0.83, SVA-Seq: 0.87).

In addition, when there are large dispersion differences across batch, ComBat-Seq is able to better control false positive rates. When applied on data with no mean batch effect and a 3-fold dispersion differences, ComBat-Seq generates the smallest FPR of 0.039, compared to the other methods (including batch as a covariate: 0.043, original ComBat on logCPM: 0.046, RUV-Seq: 0.044, SVA-Seq: 0.049). The false positive rates using data adjusted by ComBat-Seq further decrease as the level of dispersion difference increases (0.031 at the 4-fold dispersion difference, compared to the 0.039 FPR at the 3-fold difference). This suggests that ComBat-Seq is better at handling dispersion / variance batch effect in the data compared to the other available methods.

When there is no dispersion difference across batch, ComBat-Seq yields higher false positive rates with no further gain in detection power. For example, when data contain no dispersion batch effect, only a 2-fold mean batch effect, ComBat-Seq, SVA-Seq, and including batch as a covariate in the differential expression model all achieve 0.97 TPR, while ComBat-Seq has the highest FPR of 0.059 (0.047 for both the other two methods). This is consistent with the intuition for batch correction. Existing methods, such as including batch as a covariate in the differential expression methods, may be sufficient in addressing batch effect in the mean. In this case, ComBat-Seq which assumes separate dispersion across batch may be redundant and lead to higher false positives. Only when there exists dispersion / variance differences in the data are methods which specify separate dispersion parameters necessary.

### Application to the GFRN signature dataset

We applied our ComBat-Seq approach to address batch effects in a real RNA-Seq dataset designed to develop pathway signatures for breast cancer progression and treatment response (Rahman *et al.*, 2017) as described in the Methods section. Figure 5 shows the scatter plot of samples projected on the first two principle components in unadjusted data, and in data adjusted by RUV-Seq, ComBat-Seq, and using the original ComBat on logCPM. We observed a strong batch effect in the unadjusted data, which was not fully addressed by RUV-Seq. In the PCA of ComBat-Seq adjusted data, we observe the expected pattern of data if there were no batch effects, in which the control samples are clustered together, while the treated samples from three conditions are scattered at different locations. The effective adjustment of ComBat-Seq is further shown in the boxplot of proportion of explained variation by condition and batch across genes. In ComBat-Seq adjusted data, variation explained by batch is greatly reduced compared to that in the unadjusted data. These results suggest a successful adjustment of batch effect from ComBat-Seq.

**Figure 5:**
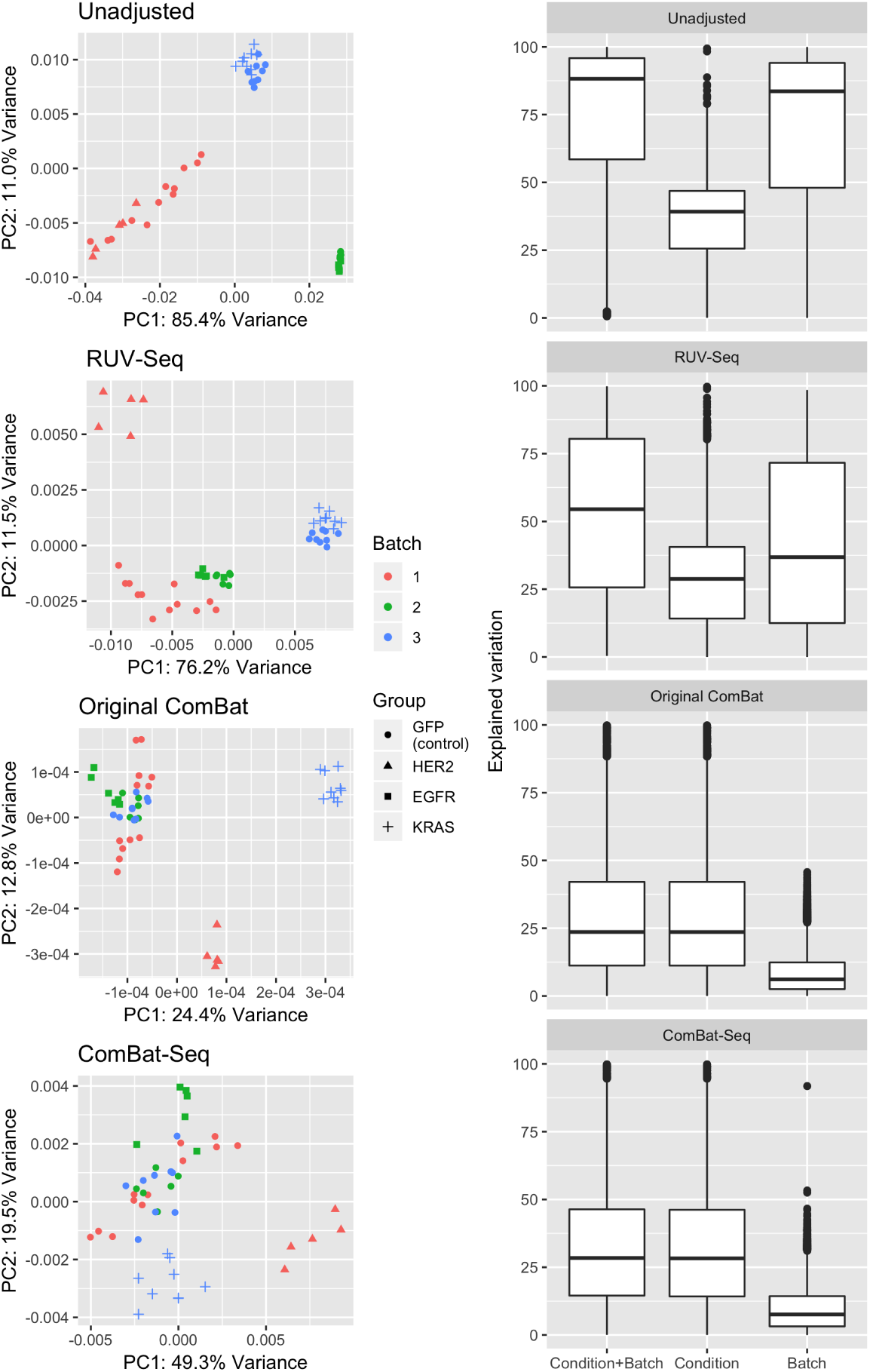
Application of ComBat-Seq for removing batch effects in a pathway activation dataset. The unadjusted data contains a strong batch effect, as samples clearly separated by batch in the principal components (top left panel, “Unadjusted”). An effective adjustment is expected to bring control samples from the 3 batches to the same level, while maintaining biological signals from the different treated samples, each of which is only present in a single batch. We observed that in the PCA plots, ComBat-Seq is able to recover the expected biological pattern, while RUV-Seq was not able to fully address the batch effect. This is further shown in the analysis of explained variation in unadjusted data, and in data adjusted by ComBat-Seq and RUV-Seq. In the ComBat-Seq adjusted data, variation explained by batch is greatly reduced compared to that in unadjusted data. Though ComBat-Seq does not show improved results in this example than the current model used on logCPM, we emphasize its benefits in increased statistical power in differential expression than the current ComBat, as we have shown in the simulation studies.

Finally, though ComBat-Seq does not show clearly improved results compared to using ComBat on logCPM in this application example, we re-emphasize that as shown in the simulations, using ComBat-Seq which preserves integer counts instead of log-transforming the data results in better statistical power in differential expression.

## Discussion and conclusions

We developed ComBat-Seq to adjust batch effects from known sources in count data from RNA-Seq studies. ComBat-Seq is able to preserve the integer nature of count data, making the analysis pipeline more compatible for RNA-Seq studies. We showed in simulations that ComBat-Seq generally out-performs other methods in terms of the impact on downstream differential expression. When variance batch effect is present in the data, ComBat-Seq is able to achieve better statistical power, while controlling false positive rates, compared to the other available methods. We further demonstrated the utility of ComBat-Seq in addressing batch effect in the GFRN signature dataset, showing its potential to recover biological signals from data affected by batch.

In simulations, we observed that when there is no true difference in dispersion across batch, applying ComBat-Seq, which specifies different dispersion parameters for batches, results in increased false positive rates compared to the other methods without further increasing the detection power. ComBat-Seq controls false positives and shows benefits in increased true positive rates only when a true dispersion batch effect is present in the data. This is consistent with the intuition of batch effect adjustment, that modifying the data in any way comes with a risk of jeopardizing biological signals in the data. Therefore, batch effects should only be adjusted when they are present and result in unfavorable impact on downstream analysis. Such observations emphasize the importance for careful diagnosis of batch effect before applying any transformation to the data.

In the simulation studies, we also compared ComBat-Seq with the commonly applied approach to transform the count matrix to logCPM, and then apply batch correction methods based on Gaussian distributions. This method essentially assumes a log-normal distribution for the data. In simulations, we observed that ComBat-Seq generally out-performs log transforming the data in terms of power and control of false positives. These results provide evidence that using appropriate probabilistic models for count data may be more beneficial than arbitrarily transforming the data.

Our study has several limitations. We used an idealistic data model in simulations, and characterized biological signals and batch effects in the form of fold changes in the average value across batch. Though there may be other methods to model count data with both condition and batch effects, our model is a valid and convenient assumption for the data, which is easy to implement with the *polyster* package. We focused primarily on addressing the unwanted impact of batch effect on downstream differential expression. It is known that batch effects may also negatively impact other biological tasks, such as developing predictive models for genomic data. Performance of batch correction in these tasks requires further evaluation, but is beyond the scope of this paper. Our ComBat-Seq method is based on a gene-wise negative binomial regression model, which, similar to other (generalized) linear models, may not work well on data with severely or even completely confounded study designs. However, batch correction in confounded designs is challenging for most if not all the state-of-the-art batch adjustment methods, and careful experimental design has been widely advised to mitigate the unfavorable impact of batch effects.

## Reproducibility

ComBat-Seq software is available at https://github.com/zhangyuqing/sva-devel. Code to reproduce the results in this paper are available at https://github.com/zhangyuqing/ComBat-seq.

## Acknowledgement

This work is supported by grants from the NIH 4P30CA006516-51 (Giovanni Parmigiani), NSF-DMS 1810829 (Giovanni Parmigiani), 5U01 CA220413-03 (W. Evan Johnson), and 5R01GM127430-02 (W. Evan Johnson). We would like to thank Dr. Solaiappan Manimaran for insightful discussions throughout the development of this method.

