## Supplementary Materials for "*ComBat-Seq*: batch effect adjustment for RNA-Seq count data"

### Parameter estimation with shrinkage

Genomic data often have small sample sizes, which makes it challenging to estimate the parameters in the ComBat-Seq model. One of the advantageous features of ComBat is the hierarchical empirical Bayes modeling, which pools information across genes for parameter estimation, making the estimates more robust for data with small sample sizes and/or outlying values (Johnson *et al.*, 2007). Similar methods for estimation with small sample size have been proposed for negative binomial regression models used on count data. The edgeR software uses a similar idea in its model, which is to estimate the dispersion parameter by maximizing a weighted likelihood, combining both gene-wise likelihood, and likelihood assuming universal dispersion across genes (Chen *et al.*, 2014).

In ComBat-Seq, we include a similar option to estimate parameters, which is inspired by the non-parametric empirical Bayes method in ComBat. In the ComBat-Seq function, users may select to use this alternative approach by setting the "shrink" parameter to TRUE (default FALSE).

The underlying methods are as follows. After fitting a gene-wise negative binomial regression model, and obtaining the batch effect parameters $(\hat{\gamma}_{gi}, \hat{\phi}_{gi})$, we adjust these parameters as a weighted average of estimates across genes:

$$\gamma_{gi}^{*} = \frac{\sum_{k \neq g} \omega_{ki} \hat{\gamma}_{ki}}{\sum_{k \neq g} \omega_{ki}} \tag{1}$$

$$\phi_{gi}^{*} = \frac{\sum_{k \neq g} \omega_{ki} \hat{\phi}_{ki}}{\sum_{k \neq g} \omega_{ki}} \tag{2}$$

where the weights are defined as

$$\omega_{ki} = L(\bar{Y}_{gi} | \hat{\gamma}_{ki}, \hat{\phi}_{ki}) = \prod_{j=1}^{n_i} d(Y_{gij} | \hat{\gamma}_{ki}, \hat{\phi}_{ki}) \tag{3}$$

$d$ represents the density function for negative binomial distributions. These formulae are directly adopted from those for the posterior estimates of mean and variance batch effect parameters in ComBat. Parameters will be estimated with the gene-wise estimates from all the other genes aside from itself. Estimates with a larger likelihood for the data will be assigned a higher weight. We then calculate the "batch-free" distribution, using the estimates as defined above. Adjusted data are generated same as before.

We evaluated this approach on the pathway signature dataset, using only the control samples from the three batches. Expected adjustment should pool all samples together. We observed that applying shrinkage to the parameter estimates tend to results in under-estimated batch effect, which leads to an under-correction of the data. Batch effects are still present in the data even after adjustment (Figure S1).

We would like to point out, however, that our proposed model is a naive extension from the ComBat non-parametric estimation approach. Deriving an empirical Bayesian approach with nega-

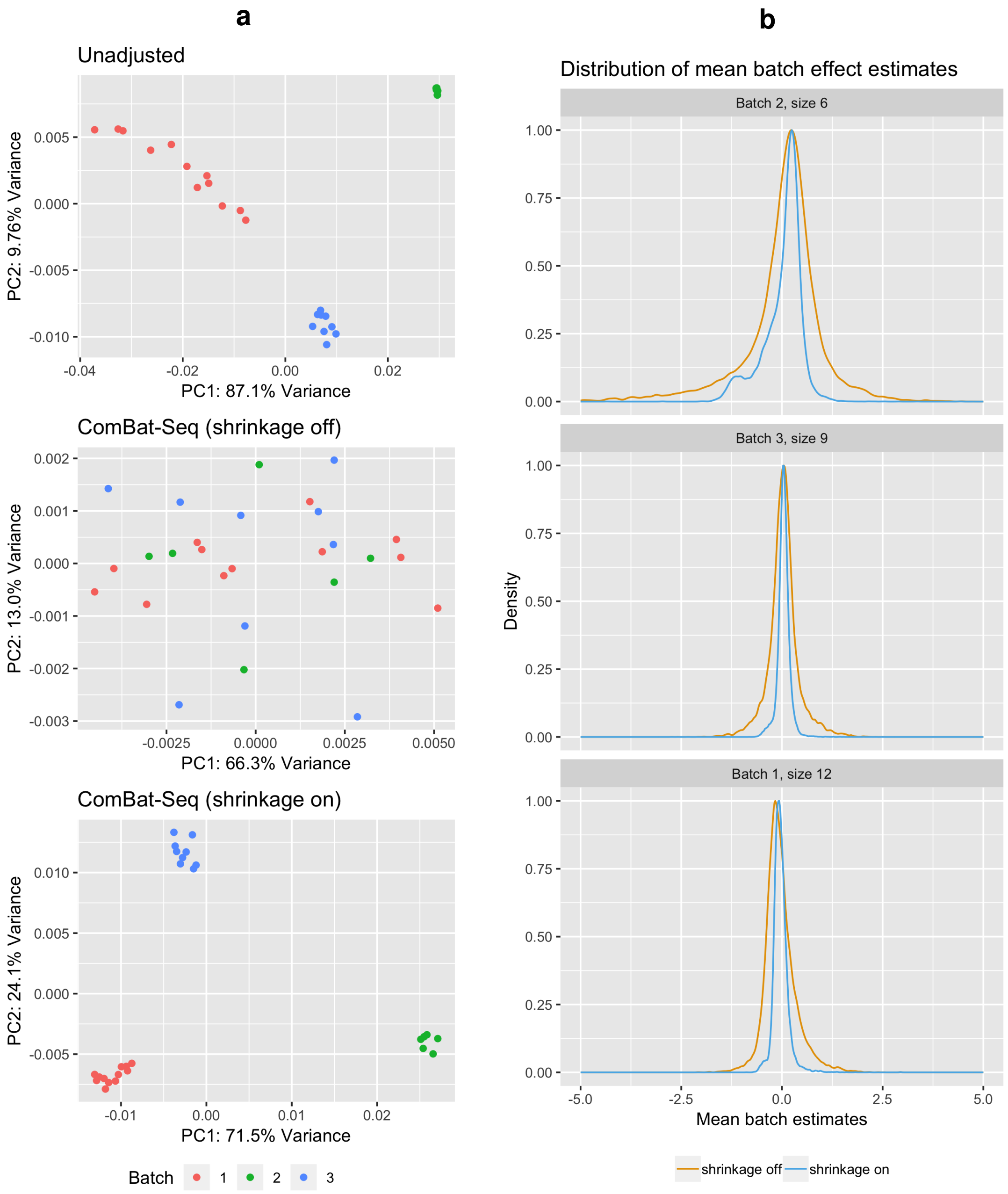


Figure S1: **Comparison between estimation with and without shrinkage.** We applied the ComBat-Seq model both with and without shrinkage on the GFRN pathway signature dataset, using only the control samples. **a)** When shrinkage is used, the estimated mean batch effect parameters of all genes tend to have a more concentrated distribution, centered at zero. This suggests that the mean batch effects are estimated to be closer to zero, an under-estimation compared to the results using ComBat-Seq without shrinkage. The under-estimation of batch effects leads to **b)** an under-correction, as shown in the PCA plots. Samples still clearly separated by batch after adjustment with the shrinkage version.

tive binomial distributed count data is challenging, for there is no conjugate distributions available. A full Bayesian approach may be feasible, but requires complicated computation and is beyond the scope of our work. We encourage further exploration on the impact of outlying counts in RNA-Seq data, and the benefits of shrinkage in batch effect estimation and adjustment.
